## Supplemental table and figures for "Linking high GC content to the repair of double strand breaks in prokaryotic genomes"

### Supplement - Linking double-strand break repair to selection for high prokaryotic GC content

October 17, 2019

S1 Table: Output of linear model relating GC content to environmental variables. The formal model was $\mathrm{GC} = \beta_0 + \beta_{\mathrm{Ku}}\mathrm{Ku} + \sum_i \beta_i \mathrm{trait}_i + \epsilon$, where GC is genomic GC content and Ku is a binary variable representing the presence/absence of Ku.

| | $\beta$ | P-value |
|---|---|---|
| (Intercept) | 0.3267217383 | 2.44868893191596E-025 |
| Ecosystem Category: Terrestrial | 0.1101532163 | 3.38243170487937E-008 |
| Ecosystem Type: Soil | 0.0333399451 | 0.1077906646 |
| Known Habitats: Soil | -0.0331866659 | 0.0551188689 |
| Habitat: Terrestrial | 0.1866541988 | 9.95695959696927E-025 |
| Nitrogen Fixation | -0.0040711787 | 0.7635124534 |
| Nitrogen production | -0.4157246148 | 3.66244214114883E-072 |
| Facultative Anaerobe | 0.0073411402 | 0.6375719405 |
| Strict Aerobe | 0.0859143088 | 7.46077194429656E-007 |
| Strict Anaerobe | -0.0143781175 | 0.4307698204 |
| Sporulation | -0.1033310344 | 3.56417677006438E-016 |
| Hyperthermophilic | -0.0834952183 | 0.0006640906 |
| Mesophilic | 0.1342814922 | 5.81638646775348E-009 |
| Thermophilic | 0.2526565354 | 1.32824551249747E-020 |
| Microaerophilic | -0.0631903302 | 3.51937747994195E-005 |
| Psychrophilic | -0.2429898729 | 7.89192914664017E-037 |
| Ku | 0.0126640175 | 0.0042992051 |

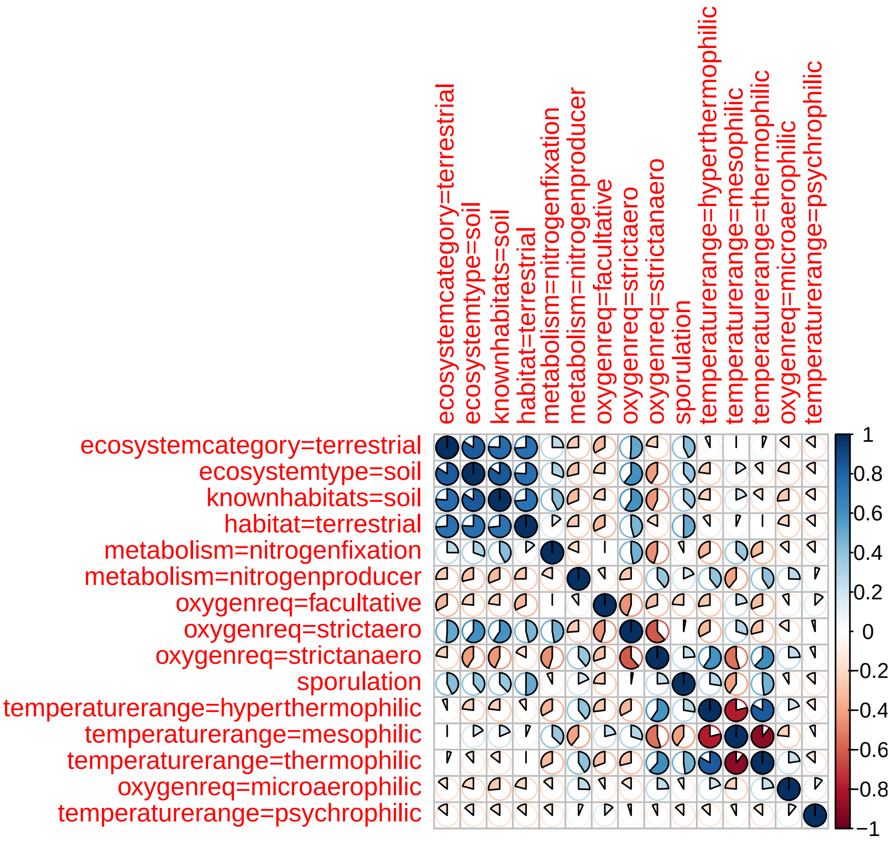


S1 Fig: The pairwise correlation between traits among species in the trait dataset. Note that some traits are highly correlated.

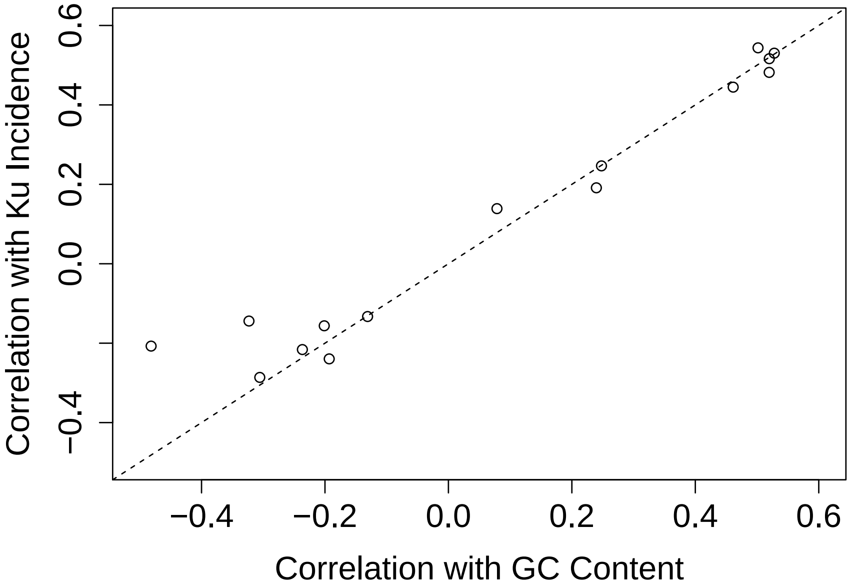


S2 Fig: The correlation of trait values for microbial species with their average genomic GC content is similar to the correlation of trait values with the presence/absence of Ku. Note that each point is an individual trait, as shown in Fig 1. The dashed diagonal line indicates the $x = y$ line. For a direct analysis of the relationship between GC content and Ku incidence among organisms see Fig 2 and Table 1.

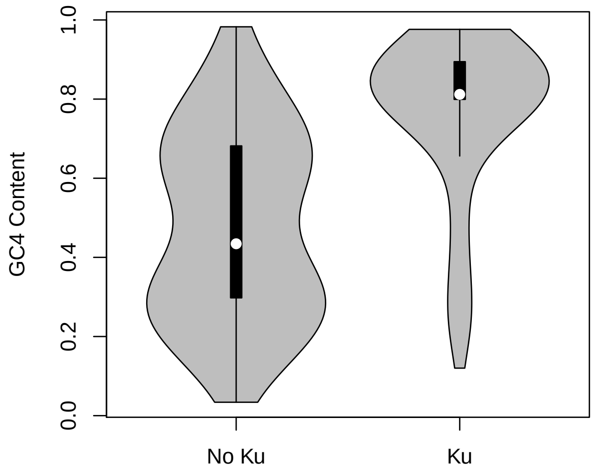


S3 Fig: GC content at fourfold degenerate sites follows a similar pattern to that of genomic GC content overall (Fig 2). The effect of Ku is significant even taking phylogeny into account using an identical approach to overall genomic GC content (Table 1).

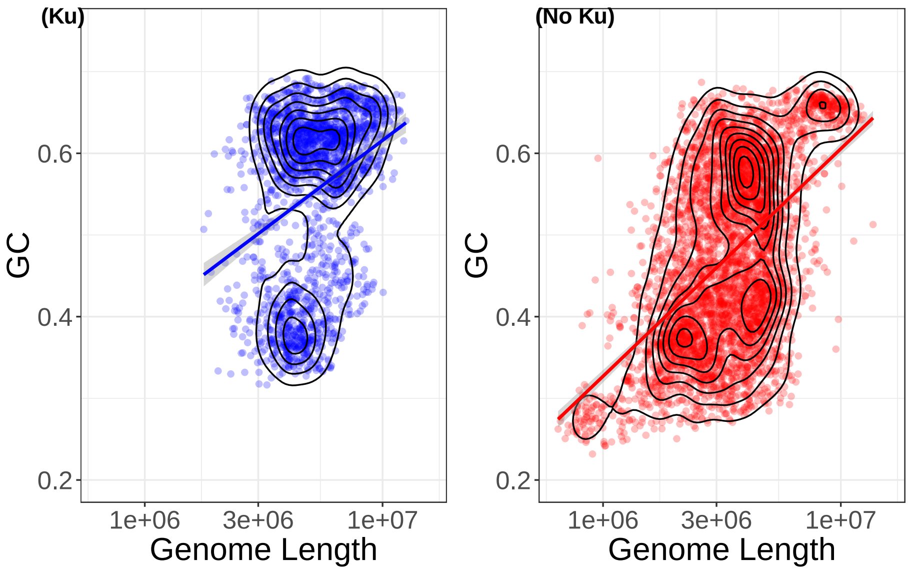


(a)

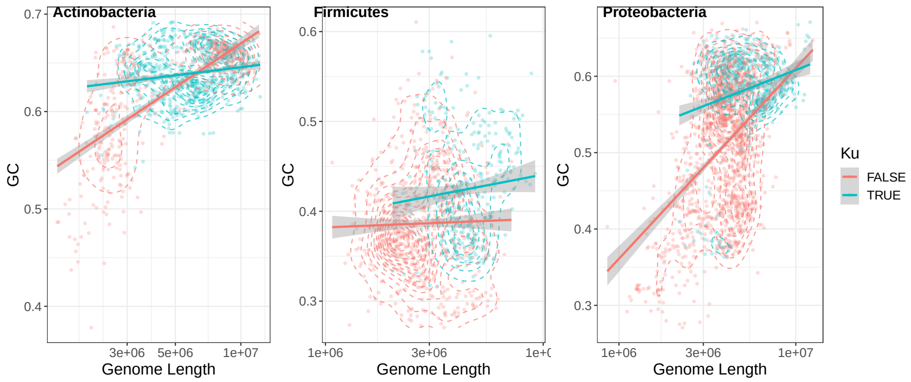


(b)

S4 Fig: While there is a positive GC content versus genome length trend, genomes with Ku have elevated GC independent of this relationship. (a) Regression and contour lines were created using default ggplot settings (b) The positive GC versus Ku relationship hold across taxa, independently of any relationship with genome length. Regressions of GC versus log genome length for Ku and non-Ku genomes shown.

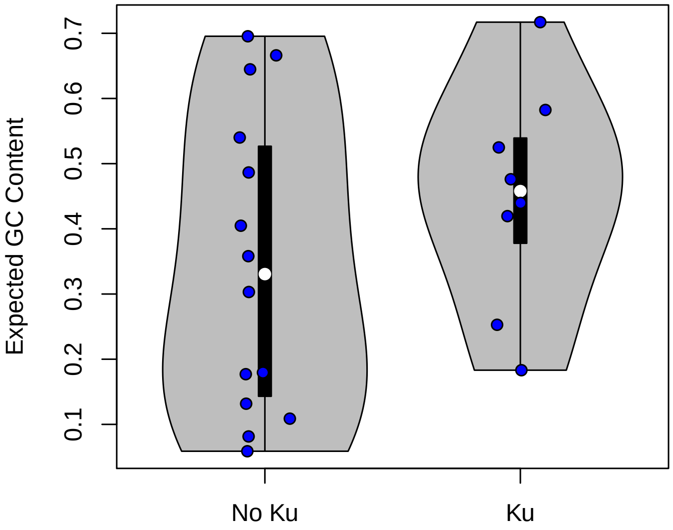


S5 Fig: Mutational bias does not appear to be associated with the NHEJ pathway. Organisms with the Ku protein did not differ significantly in their GC↔AT mutational biases from those without the Ku protein (t-test, $p > 0.34$). Estimates of mutational bias were obtained from Long et al. [1].

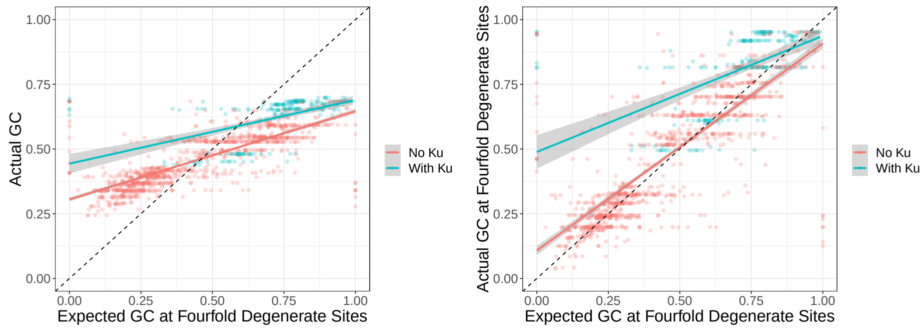


(a) (b)

S6 Fig: Genomes with Ku appear to fix GC alleles at a greater rate than expected (either due to BGC or selection). (a,b) Genomes with Ku have, on average, even greater elevation of GC over expectation than genomes without Ku. Expected GC estimated from polymorphism data; in contrast to main text Fig 3, here we only use polymorphisms at fourfold degenerate sites. This signal is conservative due to observed polymorphisms experiencing some effects of BGC/selection (see Methods for discussion).

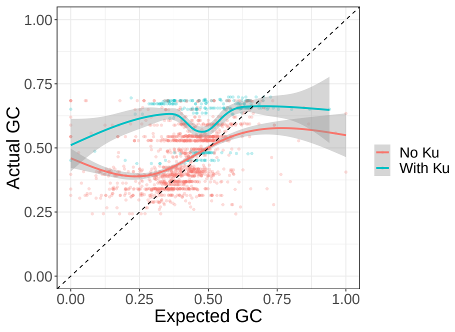


(a)

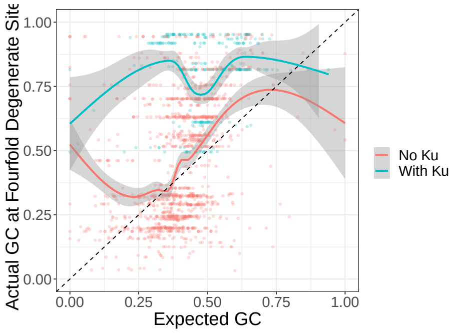


(b)

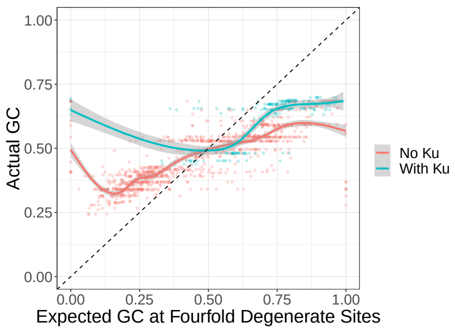


(c)

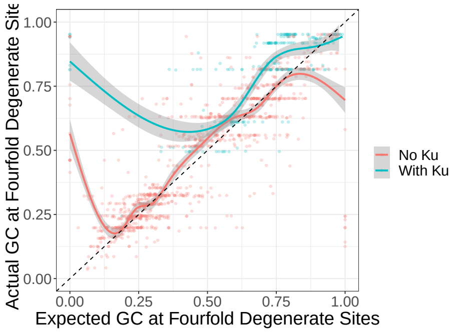


(d)

S7 Fig: Genomes with Ku appear to fix GC alleles at a greater rate than expected (either due to BGC to selection). Genomes with Ku have, on average, even greater elevation of GC over expectation than genomes without Ku. This figure is identical to panels from Fig 3 and S6 Fig except that we draw loess smoothing lines using default ggplot settings instead of linear model fits

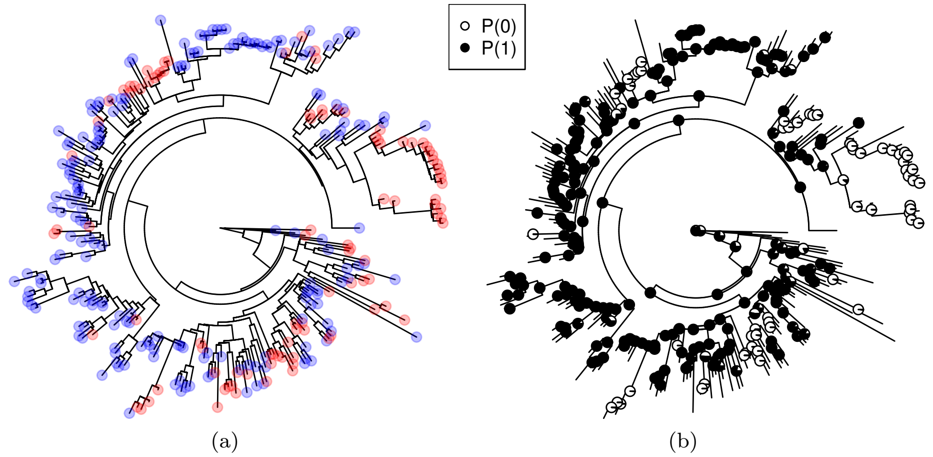


S8 Fig: Phylogeny of the *Baccilaceae* (subtree of the SILVA tree). (a) Ku presence/absence plotted on the tips of the tree as in Fig 2 (blue with, red without Ku). (b) Ancestral state reconstruction of Ku (one rate class). Each internal node is represented by a pie chart describing the probability that that organism either had (black) or did not have (white) Ku. Notice that the root and most nodes near the root are likely to have had Ku.

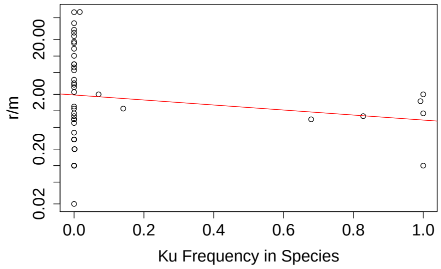


(a)

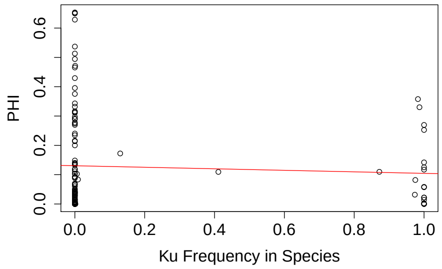


(b)

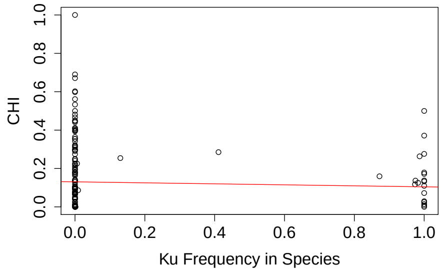


(c)

S9 Fig: Frequency of Ku presence does not appear to be positively associated with rates of homologous recombination for a species. (a) Estimated rate of recombination relative to mutation rate from Vos and Didelot [2]. (b,c) Estimated number of recombination events per gene family for species estimated with two methods by Rendueles et al. [3]. In general all of these methods give highly correlated results [3].

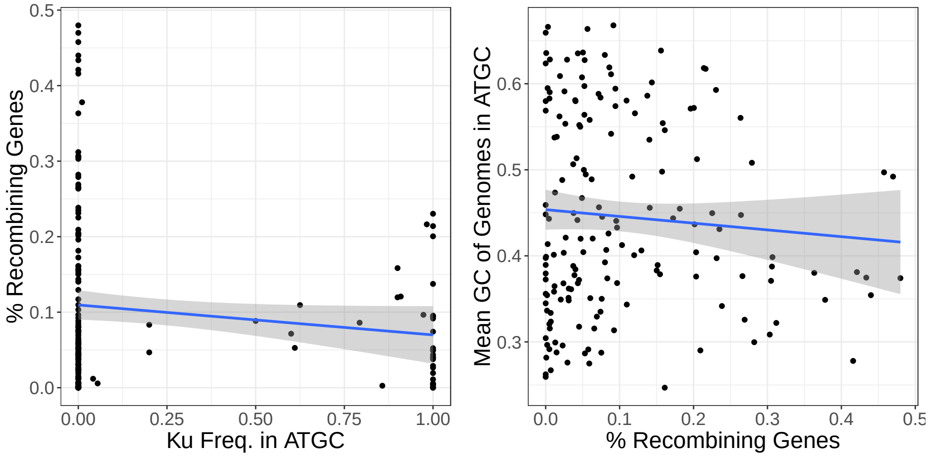


(a) (b)

S10 Fig: No relationship between genome-wide recombination frequency and (a) Ku incidence or (b) GC content in the ATGC database. We used the PHI statistic (see methods) to determine if genes within each ATGC cluster of genomes had evidence for recombination. The percent of genes with evidence for recombination (out of all genes with sufficient data to test) showed no relationship to either Ku or GC content (averaged across genomes in a particular ATGC cluster).

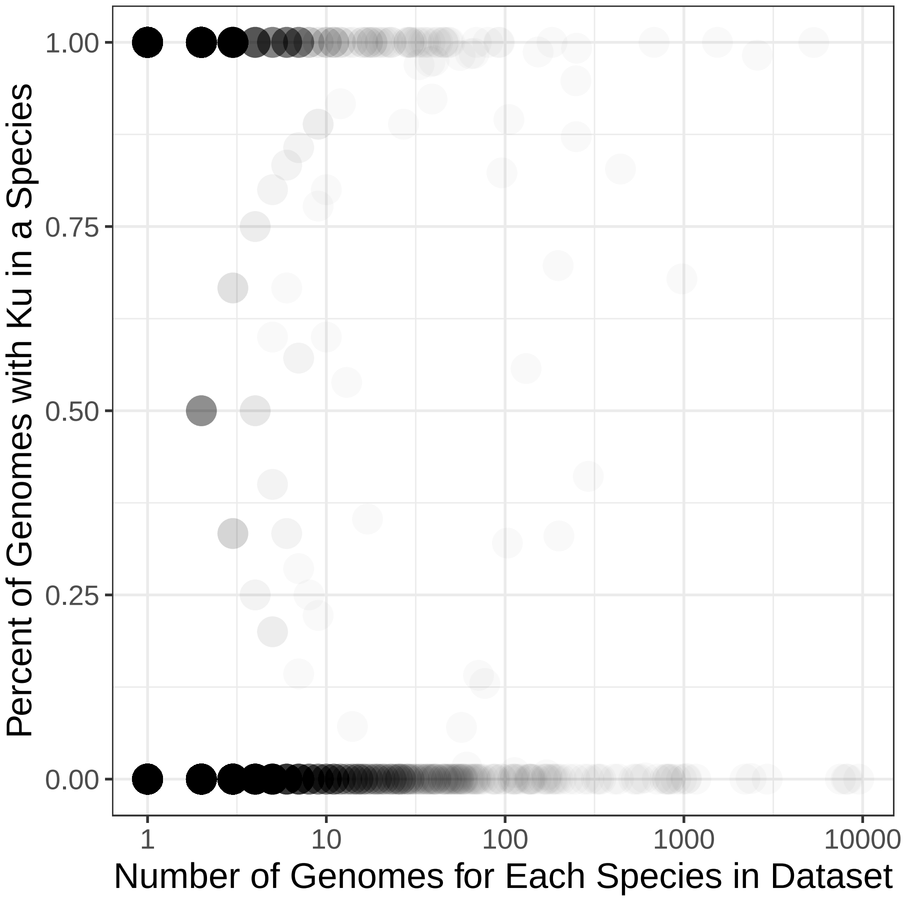


S11 Fig: Most species in RefSeq tend to always encode or always lack Ku on their genomes. Shown is the proportion of genomes within a species that have Ku (all RefSeq assemblies) plotted against the total number of assemblies in RefSeq for that species.

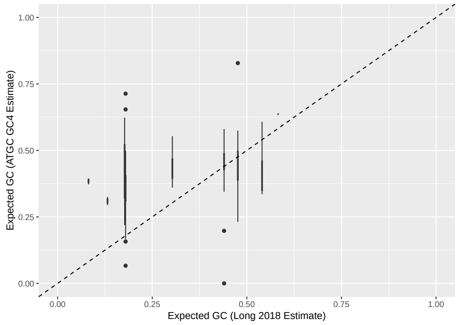


(a)

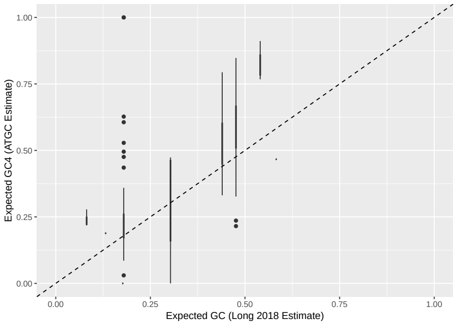


(b)

S12 Fig: We evaluate the use of polymorphisms as a proxy for mutation by comparing estimates for the few species present in both the polymorphism and mutation accumulation data. (a) Estimates based on all polymorphisms. (b) Estimates based on polymorphisms at fourfold degenerate sites. Here we see selection/BGC appears to bias the polymorphism estimates when mutation is extremely biased towards AT.
